# Natural soil suppressiveness against soilborne phytopathogens extends to the control of insect pest

**DOI:** 10.1101/2024.03.12.584584

**Authors:** Nadine Harmsen, Pilar Vesga, Gaétan Glauser, Françoise Klötzli, Clara M. Heiman, Aline Altenried, Jordan Vacheron, Daniel Muller, Yvan Moënne-Loccoz, Thomas Steinger, Christoph Keel, Daniel Garrido-Sanz

**Affiliations:** Department of Fundamental Microbiology, University of Lausanne, Lausanne, Switzerland; Institute of Earth Sciences, University of Lausanne, Lausanne, Switzerland; Centro de Biotecnología y Genómica de Plantas, Universidad Politécnica de Madrid–Instituto Nacional de Investigación y Tecnología Agraria y Alimentaria, Madrid, Spain; Neuchâtel Platform of Analytical Chemistry, University of Neuchâtel, Neuchâtel, Switzerland; Agroscope, Research Group in Entomology, Nyon, Switzerland; Université Claude Bernard Lyon 1, CNRS, INRAE, VetAgro Sup, UMR5557 Ecologie Microbienne, Villeurbanne, France

**Keywords:** benzoxazinoids, cereal leaf beetle, disease suppressive soil, phytohormones, plant-soil feedback, insect, microbiome, *Oulema melanopus*, pest control, soil suppressiveness

## Abstract

Since the 1980s, soils in a 22-km^2^ area near Lake Neuchâtel in Switzerland have been recognized for their innate ability to suppress the black root rot plant disease. Their efficacy against insect pests has not been studied. We demonstrate that natural soil suppressiveness also protects plants from the leaf-feeding pest insect *Oulema melanopus*. Plants grown in the most suppressive soil have a reduced stress response to *Oulema* feeding, reflected by dampened levels of herbivore defense-related phytohormones and benzoxazinoids, and enhanced salicylate levels in plants without the insect indicate defense-priming. The rhizosphere microbiome network of the suppressive soils was highly tolerant to the destabilizing impact of insect exposure. The presence of plant-beneficial bacteria in the suppressive soils along with priming conferred plant resistance to the insect pest, manifesting also in the onset of insect microbiome dysbiosis. This intricate soil-plant-insect feedback extends natural soil suppressiveness from soilborne diseases to insect pests.

## Introduction

Pests and pathogens cause ∼20% of global losses in major crops^1^. Among insect pests, the cereal leaf beetle *Oulema melanopus* (Coleoptera: *Chrysomelidae*) feeds on numerous species of wild and cultivated grasses, including the top-five crop wheat^2,3^, and it is considered a major threat in Europe, Asia, and North America. Losses caused by *O. melanopus* will increase as temperature rises due to climate change will expand its geographic range and amplify its impact on crops^4^. Current control practices of *O. melanopus* mainly rely on the use of chemical insecticides^5^, although biological control by endoparasitoids and entomopathogenic nematodes has long been reported^6,7^. Nonetheless, the existing pressure for reducing the use of chemical insecticides, driven by their harmful impact on the environment, pushes the development of environmentally friendly and sustainable approaches for pest mitigation.

Natural soil suppressiveness has long been observed and refers to the capacity of certain soils to confer plant protection against diseases caused by specific phytopathogenic fungi, oomycetes, bacteria or nematodes^8–15^. In certain cases, soil suppressiveness depends on a few key protective microbial populations present in the soil^9,16–21^. One of the best-documented cases is the natural suppressiveness of Swiss soils near Lake Neuchâtel against the black root rot pathogen *Thielaviopsis basicola*^11,21,22^. These soils were first studied in the 1980’s, and a relationship was found between their disease suppressiveness, physicochemical characteristics and microbiome composition^11,22,23^. More specifically, soils containing vermiculite clay supported the proliferation and activity of pathogen-inhibiting *Pseudomonas* species, contrary to disease-favoring (so-called conducive) soils containing illite clay^11,21,22,24^. These *Pseudomonas* release antifungal compounds that effectively antagonize *T. basicola* in soil, protecting tobacco crops^11,25^. Several representatives of *Pseudomonas* can also colonize and kill pest insects with insecticidal toxins^26–29^. Therefore, bacteria present in naturally suppressive Swiss soils could potentially protect plants against both fungal and insect pests. Although augmentation of entomopathogen populations or microbiome management have been proposed as strategies to mitigate pest incidence in soils^30,31^, natural soil suppressiveness towards pest insects has not been documented.

The microbiome of *O. melanopus* consists of endosymbionts such as *Wolbachia*, and other bacteria that can be significantly influenced by the host plant^32^, which can itself be influenced by the soil microbiome, for example through plant immune response priming against microbial and insect attacks^33,34^. This is known to be mediated by phytohormones such as jasmonic acid and salicylic acid^35–37^, or by cyanogenic compounds, such as linamarin and lotaustralin^38^. In the grass family *Poaceae*, benzoxazinoids (BXs) are also a predominant class of plant defense molecules with antimicrobial and insecticidal activity^39–43^. BXs are secondary metabolites that are stored in a stable glucoside form in vacuoles within plant cells, released upon cell damage, and transformed into the active, unstable aglucone form^39,40,44^. The most abundant BX in wheat is DIMBOA, which has a high toxicity against some insects ^40,44–46^. The soil microbiome can also influence the plant defense metabolism, priming it against leaf-chewing pest insects^47–50^. Thus, monitoring the concentrations of these three metabolite classes can provide insights into how natural soil suppressiveness can influence the plant defense responses.

In this work, we tested whether disease-suppressive soils can also protect from insect pests, by studying the interaction of *O. melanopus* with wheat in four soils with contrasting levels of suppressiveness against *T. basicola*. We assessed changes in insect larval mortality and herbivory, plant defense metabolites, as well as the microbiome composition within the soil, rhizosphere, leaves and insect larvae associated with plants grown in the different soils. Our results demonstrate that soils manifesting natural suppressiveness against soilborne plant pathogens can also offer protection against herbivorous insects through a complex soil-plant feedback involving the presence of plant-beneficial bacteria and the priming of systemic plant defense responses.

## Results

### Disease suppressiveness remains in soils after decades

We collected six soils from the Lake Neuchâtel region in Switzerland (**Fig. 1a**) that have been studied in the past decades for their natural suppressiveness against the tobacco black root rot caused by the soilborne fungal pathogen *T. basicola*^11,21–23,51^. We reassessed the ability of the soils to naturally suppress *T. basicola* infection in tobacco plants. We found that the soils previously classified as conducive (C6, C10, and C112) retained this characteristic as there was a significantly higher disease severity (60-100%) in tobacco plants growing in the *T. basicola*-inoculated soils compared to the non-inoculated soils (**Fig. 1b**). In contrast, the three soils previously identified as suppressive soils^23,51^ (S16, S7 and S8) produced differing results. Tobacco plants growing in the soil S16 exhibited significantly lower disease severity (<25%) when inoculated with *T. basicola* compared to the conducive soils. The soil S7 presented intermediate disease suppressiveness (**Fig. 1b**), with a range of 25-75% disease severity that differed significantly from S16 and the conducive soils, in agreement with previous findings^23,51,52^. Plants in the uninoculated S8 soil showed similar disease severity to the inoculated conducive soils. The long-standing suppressive or conducive status was confirmed in five of the six soils investigated, and four soils, C10, C112, S7, and S16 were selected for further analysis.

**Fig. 1.**
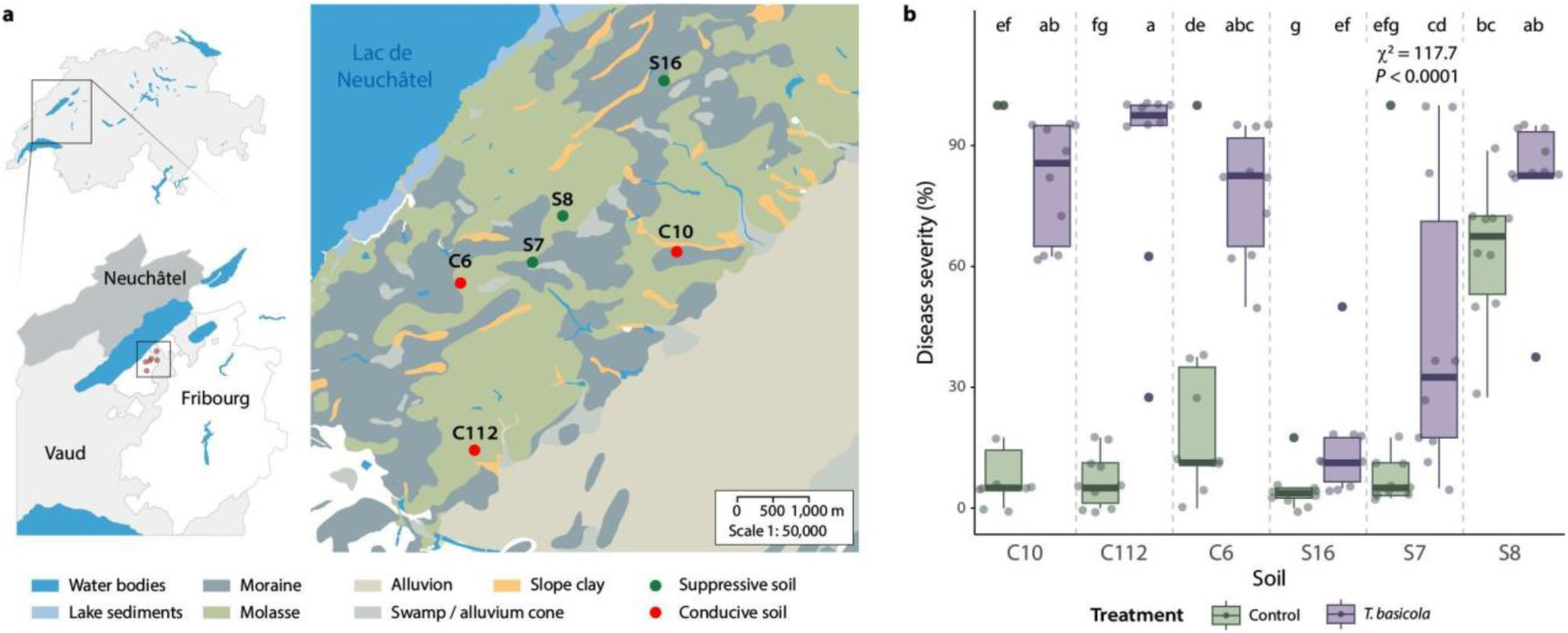
Geographical location of the soils and natural suppressiveness against *Thielaviopsis basicola*. **a**, Soil collection sites within the Swiss Lake Neuchâtel region and map (right) emphasizing the key lithological components. Conducive/Suppressive identity of soils according to previous reports. **b**, Tobacco black root rot suppressiveness of the six soils tested in this study towards the fungal pathogen *T. basicola*. Boxplots represent the disease severity in plants grown in the six soils inoculated with *T. basicola* (purple) or left uninoculated (green). The disease severity was assessed by scoring the percentage of the root affected by the pathogen. Points represent individual replicates. Ten replicates per soil and condition were performed. Differences were assessed using the Kruskal-Wallis test and post hoc analysis using Fisher’s least significant difference (LSD). The *P* values were corrected using the false discovery rate (FDR). Different letters indicate significant differences between groups (*P* ≤ 0.05). The maps were generated using data from the Swiss Federal Office of Topography swisstopo, where the original map was sourced and vectorized.

### Soil suppressiveness increases insect mortality and reduces leaf damage

We then evaluated the effect of conducive or suppressive soils towards larvae of the cereal leaf beetle *O. melanopus*. Larvae were left for 11 days on wheat plants that were grown in the four selected soils and their survival was assessed daily (**Fig. 2a**). There was a significant difference between the C10 conducive soil, which produced the fewest larvae deaths, and the S16 suppressive soil, where all larvae had died by day nine (Cox model, *P* = 0.0003, **Fig. 2b, Supplementary Note 1**). The soils C112 and S7 did not show any statistical difference compared to the other soils. Additionally, the *O. melanopus* larval mortality at the midpoint, defined by the time of the first and last evaluated deaths, and at the endpoint, was used to calculate the percentage of dead and alive larvae. The soil S16 had the highest percentage of dead insect larvae at both time points (90-100%), while C10 had the lowest (35-65%, **Fig. 2b**). Between the mid- and endpoint, C112 could be seen to support an increase in mortality compared to S7, potentially backing the notion that S7 is a partially suppressive soil.

**Fig. 2.**
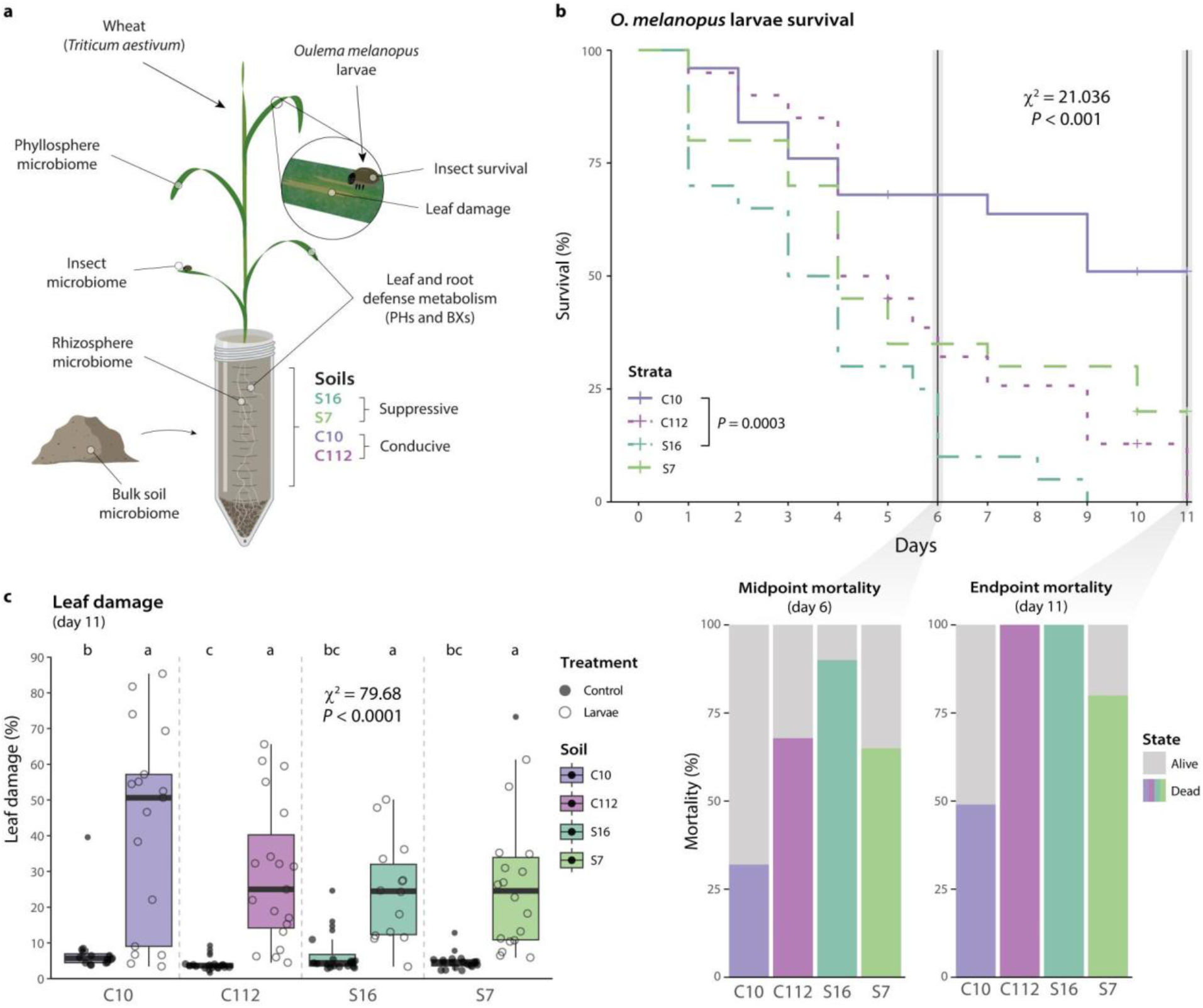
Survival of *Oulema melanopus* larvae and leaf damage on wheat plants grown in the four soils. **a**, Experimental set-up followed in this study. PHs, phytohormones; BXs, benzoxazinoids. **b**, Kaplan-Meier survival curves showing insect larvae survival over 11 days (*n* = 20, two experimental runs per condition). The vertical dark-gray lines indicate the midpoint (day 6) between the first insect death in the experiment and the endpoint of the experiment (day 11). Statistical differences were assessed using the mixed-effects Cox proportional hazards model and Tukey’s post hoc test. The *P* values were corrected by FDR. Barplots below show the percentage of mortality at the midpoint (left) and the endpoint (right) of the experiment. Colored portions represent the percentage of dead larvae. **c**, Boxplots representing the percentage of damaged leaf surface area on plants with and without *O. melanopus* larvae in the four soils at the end of the experiment (day 11). The points represent individual replicates (*n* ≥ 15, two experimental runs). Statistical differences were assessed using the Kruskal-Wallis test and LSD post hoc analysis. The *P* values were corrected by FDR. Different letters indicate significant differences between groups (*P* ≤ 0.05).

The leaf area damaged by feeding *O. melanopus* larvae showed a strong difference between groups with and without larvae (**Fig. 2c**), with an average of ∼30-60% of damaged leaf area observed in the plants exposed to herbivory, compared to less than 10% in the controls. We also observed a lower mean percentage of damaged leaf area in the suppressive soils (<30%) compared to the conducive soils.

### Salicylate-mediated priming in leaves reduces plant stress in suppressive soils

To evaluate the plant response to the presence of *O. melanopus* larvae in the four soils, we analyzed the concentration of five defense-related phytohormones^53–55^ in leaves and roots: abscisic acid (ABA), jasmonic acid (JA), jasmonyl-isoleucine (JA-Ile), 12-oxo-phytodienoic acid (OPDA), and salicylic acid (SA, **Fig. 3, Extended Data Fig. 1**). The presence of the herbivorous larvae significantly increased the concentration of all the assessed phytohormones in the leaves of plants from all soils (**Fig. 3a**). In contrast, except for ABA, the presence of insect larvae did not significantly impact phytohormone concentrations in the roots (**Extended Data Fig. 1**).

**Fig. 3.**
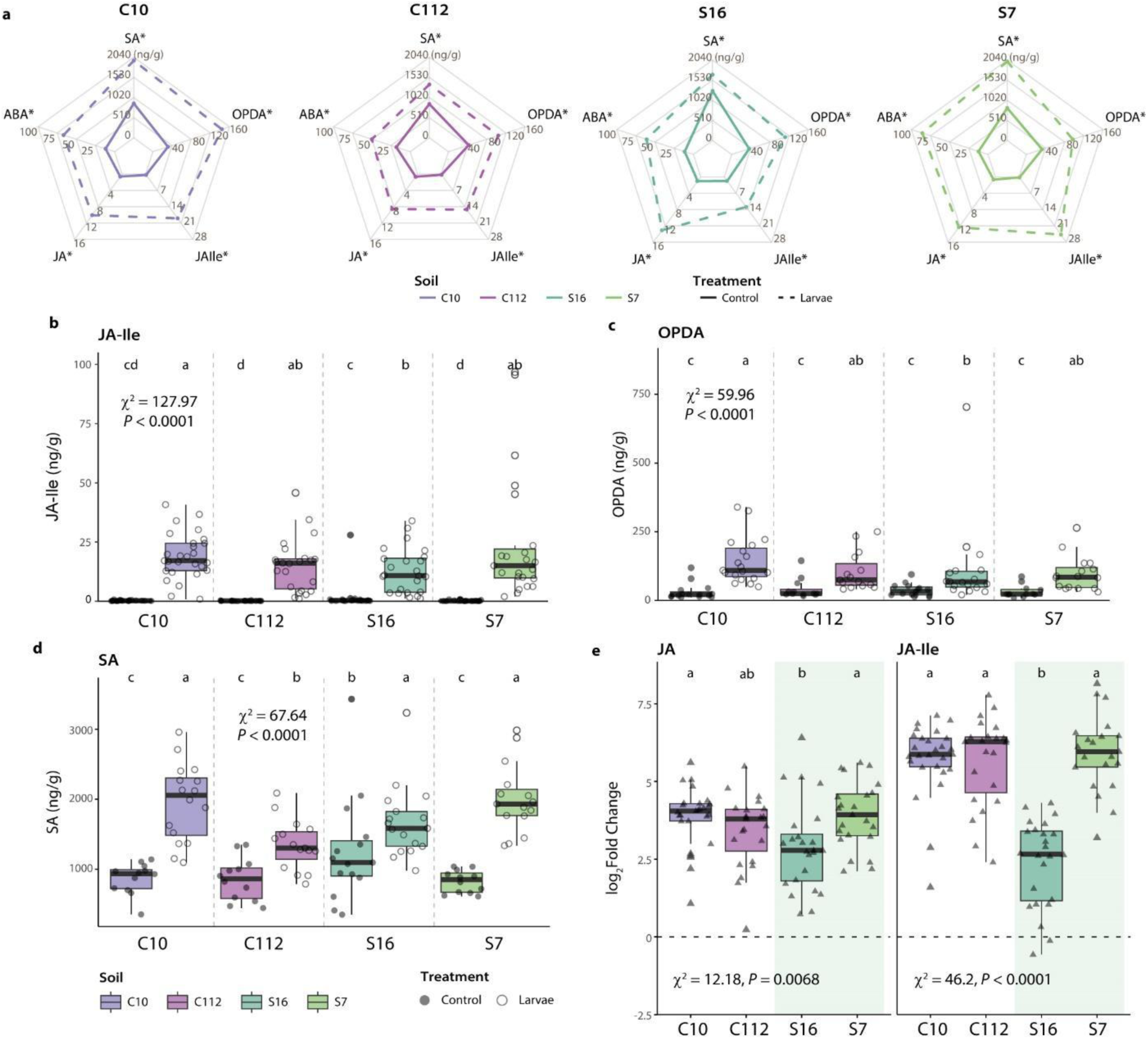
Changes in levels of stress-related phytohormones in wheat leaves exposed to *Oulema melanopus* larvae. **A**, Spider plots show insect and control group concentrations of abscisic acid (ABA), jasmonic acid (JA), jasmonyl-isoleucine (JA-Ile), 12-oxo-phytodienoic acid (OPDA), and salicylic acid (SA) in leaf samples in the soils C10, C112, S16, and S7. Asterisks denote phytohormones whose concentrations differed significantly between insect and control groups (Kruskal-Wallis, *P* ≤ 0.05). For root samples, see **Extended Data Fig. 1a. b-d**, Boxplots representing the concentration of (**b**) JA-Ile, (**c**) OPDA, and (**d**) SA in leaf samples of plants that had been exposed or not to the insect in the four soils (*n* ≥ 18 for JA-Ile, *n* ≥ 12 for OPDA and SA, at least two experimental runs were performed). The points represent individual replicates. **E**, Log_2_ fold change of JA (left) and JA-Ile (right). Fold change was calculated by dividing the concentration in insect-exposed plants by the mean concentration in the control group. Values above zero signify an increase in concentration with respect to the control. Triangles represent individual replicates. Statistical differences were assessed using the Kruskal-Wallis test and LSD post hoc analysis. The *P* values were corrected by FDR. Different letters indicate significant differences between groups (*P* ≤ 0.05). For details, see **Extended Data Fig. 1**.

Concentrations of the jasmonate precursor OPDA, and of JA-Ile, both involved in plant defense signaling when faced with insect herbivory^35–37,56^, were highest in the conducive C10 soil and lowest in the highly suppressive S16 in the presence of *O. melanopus* larvae (**Fig. 3bc**). Similarly, changes in JA and JA-Ile levels induced in wheat leaves upon insect exposure were smaller in S16 soil than in the other soils (**Fig. 3e**), meaning that plants were less stressed. The S16 control group also had a significantly higher level of salicylic acid (SA) in leaves compared to the other control groups, indicating that the plants were primed against biological threats in this soil (**Fig. 3d**).

### Dampened levels of insect defense-related benzoxazinoids in suppressive soils

The concentrations of six benzoxazinoids (BXs) involved in plant responses to insect damage^40,44,45^, including the more active DIMBOA and its storage form DIMBOA-Glc, were also analyzed in leaf and root samples (**Extended Data Fig. 2**, **Supplementary Note 2**). Additionally, two cyanogenic glycosides, linamarin and lotaustralin were measured (**Extended Data Fig. 3**). Unlike the phytohormones, the concentration of most BXs in leaf samples was not affected by *O. melanopus* larvae regardless of the soil used. Exceptions to this were DIMBOA, HBOA-Glc, and HMBOA-Glc in the soils C10 and C112, DIMBOA-Glc in S16, and HMBOA-Glc in S7 (**Fig. 4a**). Plant leaves exposed to *O. melanopus* larvae in the suppressive S16 and S7 soils showed a trend of lower DIMBOA concentration compared to those in the conducive soils, which could not be observed for the storage form DIMBOA-Glc (**Fig. 4bc**). Similarly, changes in DIMBOA levels upon herbivory exposure were significantly lower in the suppressive S16 and S7 soils compared to the conducive C10 and C112 soils, where DIMBOA concentrations almost doubled in the presence of larvae (**Fig. 4d**). While in the absence of insects the ratio of DIMBOA:DIMBOA-Glc in the leaves was similar, the concentration of DIMBOA relative to DIMBOA-Glc increased in the insect-exposed groups in all soils except in the highly suppressive S16 soil (**Fig. 4e**). The cyanogenic glycosides did not differ significantly among any condition or soil type (**Extended Data Fig. 3**).

**Fig. 4.**
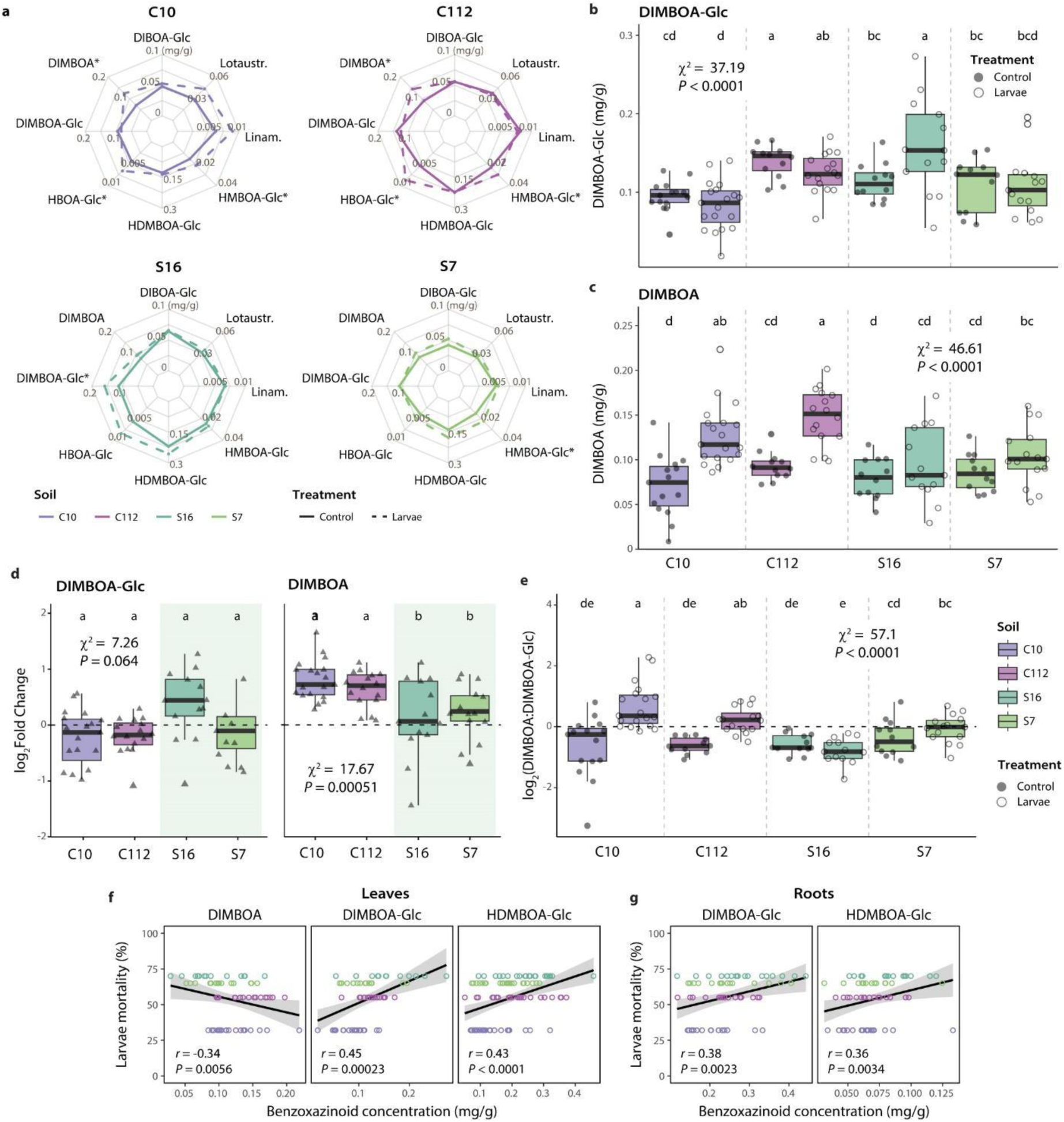
Changes in benzoxazinoid (BX) concentration in wheat leaves exposed to *Oulema melanopus* larvae. **a**, Spider plots show *O. melanopus* larvae and control group concentrations of DIBOA-Glc, DIMBOA, DIMBOA-Glc, HDMBOA-Glc, HBOA-Glc, and HMBOA-Glc, and two cyanogenic compounds (lotaustralin: lotaustr., linamarin: linam.) in leaf samples from the soils C10, C112, S16, and S7. Asterisks denote BXs whose concentrations differed significantly between insect and control groups (Kruskal-Wallis, *P* ≤ 0.05). For root samples, see **Extended Data Fig. 2a. b,c**, Boxplots representing the concentration of DIMBOA-Glc (**b**, *n* ≥ 12) and DIMBOA (**c**, *n* ≥ 12) in leaf samples exposed or not to the larvae in the four soils. Two experimental runs were performed. **d**, Log_2_ fold change of DIMBOA-Glc (left) and DIMBOA (right). Fold change was calculated by dividing the concentration in treated individuals by the mean concentration in the control group. Values above zero signify an increase in concentration over the control. **e**, Boxplot of the log_2_ DIMBOA:DIMBOA-Glc ratio in leaf samples exposed or not to *O. melanopus* larvae in the four soils. Values above zero signify a higher concentration of DIMBOA. Statistical differences were assessed using the Kruskal-Wallis test and LSD post hoc analysis. The *P* values were corrected by FDR. Different letters indicate significant differences between groups (*P* ≤ 0.05). For details, see **Extended Data Fig. 2. f,g**, Significant Spearman correlations between individual benzoxazinoid concentrations and larval mortality at day 5 in (**f**) leaves or (**g**) roots.

We examined the relationship between BXs and mortality of *O. melanopus* larvae on day 5 (same day of BXs extraction). The concentration of DIMBOA-Glc and HDMBOA-Glc in leaves and roots showed a significant positive correlation with insect mortality (Spearman, *r* > 0.36, *P* < 0.01, **Fig. 4fg**), while a negative correlation was observed for DIMBOA in leaves (Spearman, *r* = -0.34, *P* = 0.0056).

### Onset of microbiome dysbiosis in insects feeding on suppressive soil-grown plants

The microbiomes of the four soils, the rhizosphere and phyllosphere of the wheat plants growing in them, and *O. melanopus* larvae feeding on these plants were analyzed. There were systematically more amplicon sequence variants (ASVs) and higher alpha diversity in the soil and rhizosphere microbiomes than in the wheat leaves or in the *O. melanopus* larvae microbiomes (**Extended Data Fig. 4)**. Significant differences in the larval microbiomes were found between soils based on Bray-Curtis dissimilarities, with S7 and C112 clustering together and distant from S16 and C10 (**Fig 5a**). Differential abundance analysis of ASVs between *O. melanopus* larvae feeding on plants growing in the C10 and S16 soils revealed distinct *Pantoea, Wolbachia,* and endosymbiont (*Enterobacterales*) populations (**Fig. 5b**), which contribute more than ∼75% of the larval microbiomes (**Fig. 5c)**. No significant differences were found in the relative abundance of these three genera in *O. melanopus* larvae based on soil type, except for more abundant endosymbionts in larvae on plants grown in the C112 and S7 soils (**Fig. 5cd**). However, a lower median relative abundance of *Wolbachia* and other endosymbionts in the most suppressive S16 soil coincided with higher *Pantoea* abundance, reflecting individual ASV changes (**Fig. 5e**), and the onset of microbiome dysbiosis.

**Fig. 5.**
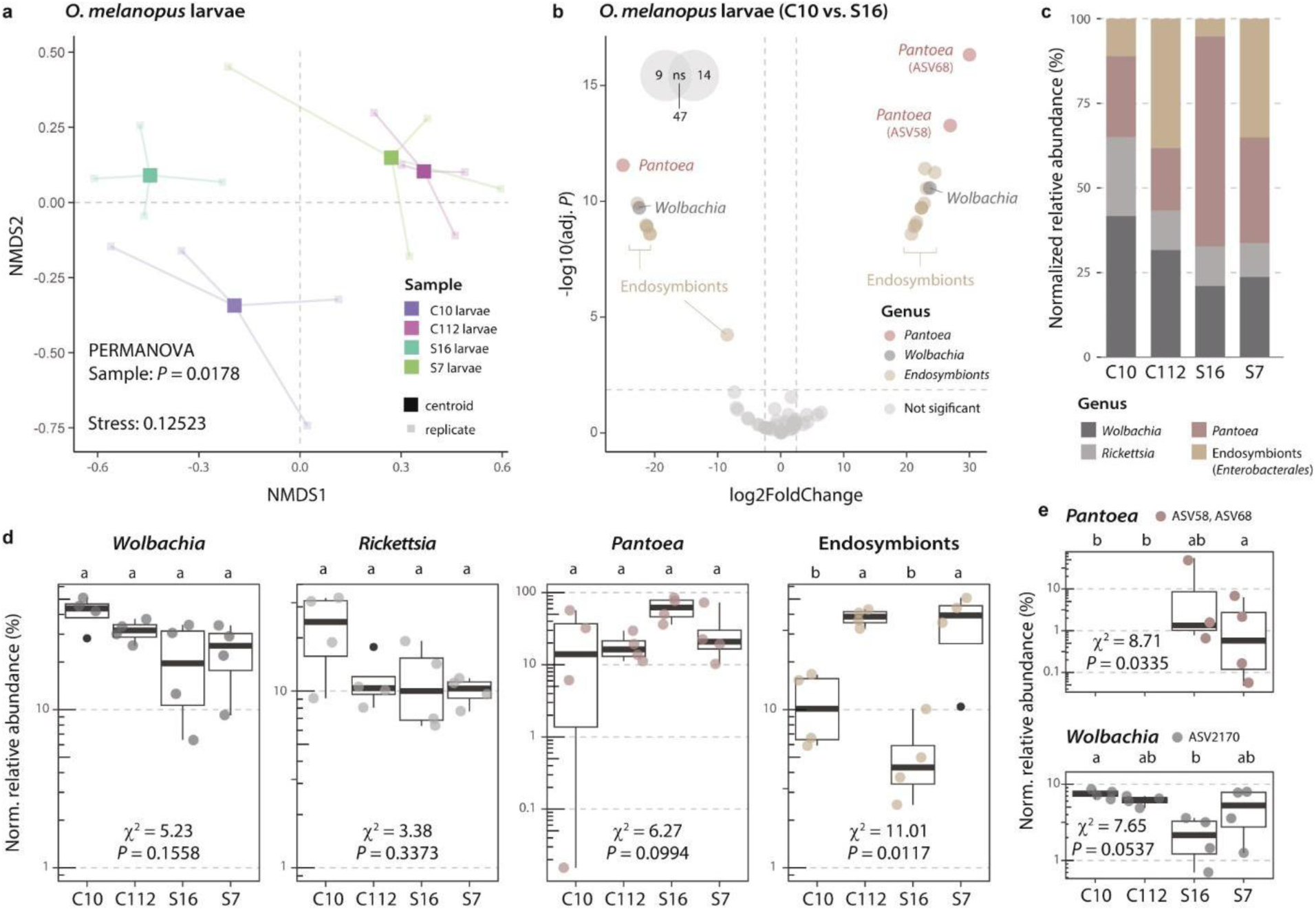
Microbiome composition of *Oulema melanopus* larvae in response to different levels of soil suppressiveness. **a**, Non-metric multidimensional scaling (NMDS) ordination analysis of microbiomes of *O. melanopus* larvae feeding on wheat plants grown in soils C10, C112, S16, and S7. Statistical differences between samples were assessed using PERMANOVA. **b**, Differential abundance analysis of amplicon sequence variants (ASVs) between the microbiomes of larvae feeding on plants grown in C10 soil (left) compared to those from the soil S16 group (right). Significantly abundant ASVs (colored dots) were considered with an adjusted (adj.) *P* ≤ 0.01 and a |log2FoldChange| ≥ 2.5. **c**, Cumulative sum scaling (CSS)-normalized relative abundance of the top 50 ASVs present in *O. melanopus* larvae. Abundances represent mean values (*n* = 4). **d,e**, Individual differences in (**d**) genera relative normalized (Norm.) abundance or (**e**) specific ASVs (*n* = 4). ASV58 and ASV68 are shown together, as they only differed in a single nucleotide. Significant differences between relative abundances were assessed using the Kruskal-Wallis rank sum test and LSD post hoc analysis. The *P* values were corrected by FDR. Different letters indicate significant differences between groups (*P* ≤ 0.05).

### Rhizosphere microbiome in suppressive soil resists to pest-induced destabilization

The microbiomes of the four soils were significantly different based on Bray-Curtis dissimilarities (PERMANOVA, *P* = 0.0001), with the largest difference between S16 and C10 (PERMANOVA, *P* = 0.0072) (**Fig. 6a**). Leaf-feeding by *O. melanopus* larvae did not have a significant effect on the overall composition of the wheat rhizosphere microbiomes in any of the soils (**Fig. 6a**). Differential abundance analysis of the ASVs in the C10 and S16 rhizospheres of plants exposed to *O. melanopus* larvae revealed populations of several bacterial genera, including *Pseudomonas*, that were found enriched in one soil or the other (**Fig. 6b**). The presence of larvae also coincided with an enrichment of certain *Pseudomonas* ASV populations in the C10 and S16 rhizospheres (**Fig. 6c**). An exact ASV sequence match for *Pseudomonas protegens* was detected in the S16 rhizosphere, with no differences between the larvae and control groups. This ASV remained undetectable in any of the other soils (**Fig. 6d**). Additionally, there were distinct patterns in rhizosphere-associated genera between the different soils. For example, *Herbaspirillum* and *Rahnella* were not detected in the C10 soil, while *Streptomyces* was more abundant in this soil (**Fig. 6e, Extended Data Fig. 5**).

**Fig. 6.**
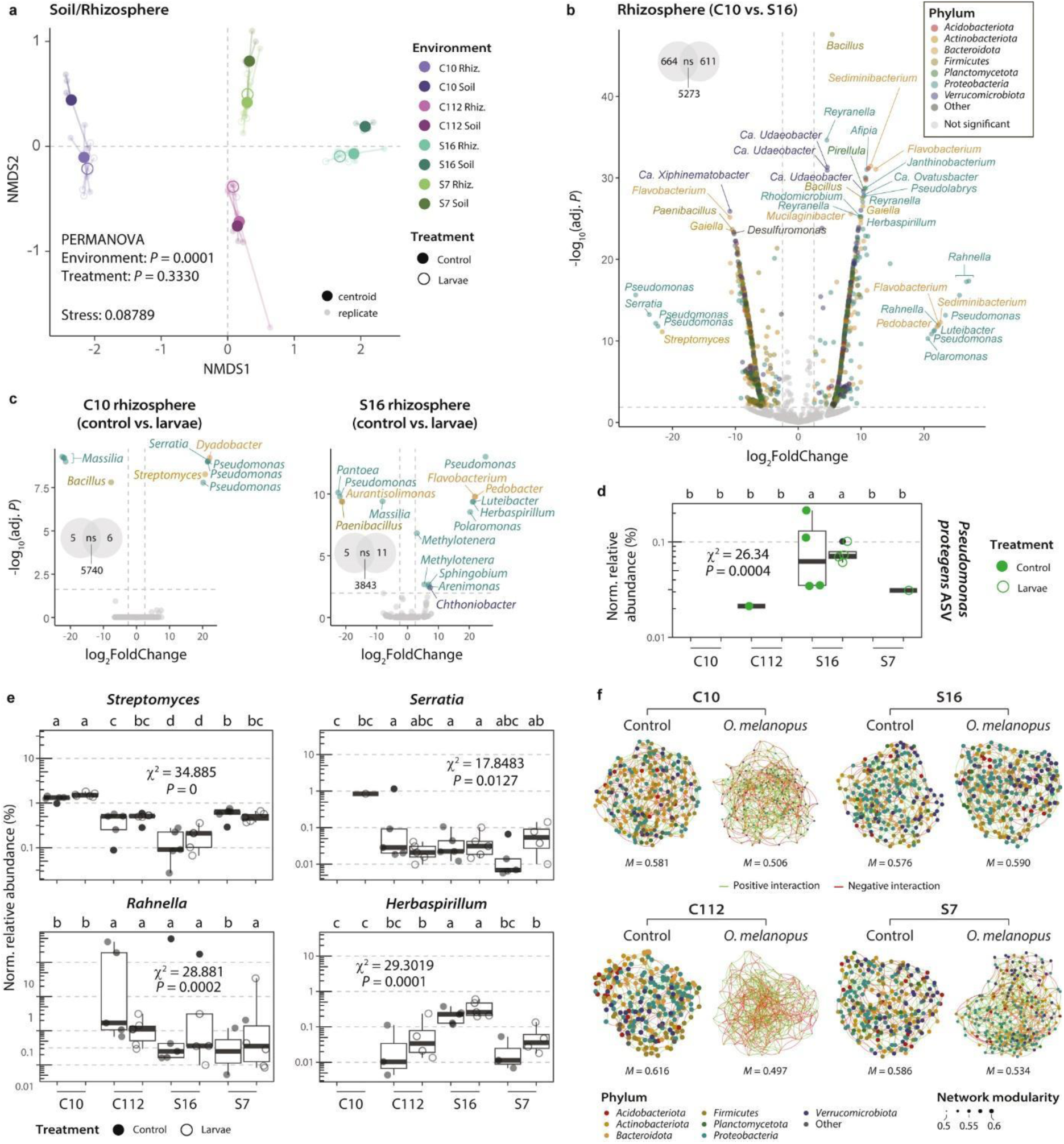
Effect of different soil suppressiveness levels on rhizosphere microbiome composition. **a**, Non-metric multidimensional scaling ordination analyses of soil and rhizosphere (Rhiz.) samples. Statistical differences between environments and treatments were assessed using PERMANOVA. **b,c**, Differential abundance analyses between conditions (C10 in relation to S16 in **a**, or control in relation to larvae in **c**). Significantly abundant ASVs (colored dots) were considered with an adjusted (adj.) *P* ≤ 0.01 and a |log2FoldChange| ≥ 2.5. **d,e**, CSS-normalized (Norm.) relative abundance of (**d**) *Pseudomonas protegens* CHA0^T^ (exact ASV sequence match) or (**e**) different genera in the rhizospheres of plants grown in the four soils exposed or not to *Oulema melanopus* larvae (*n* ≥ 4, values = 0 have been omitted from the representation, see **Extended Data Fig. 5**). Significant differences were assessed using the Kruskal-Wallis rank sum test and LSD post hoc analysis. The *P* values were corrected by FDR. Different letters indicate significant differences between groups (*P* ≤ 0.05). **f**, Sparse inverse covariance networks among the top 250 ASVs (colored dots) from rhizosphere microbiome samples in control plants or those exposed to *O. melanopus* herbivory. Edges represent positive and negative interactions in green and red, respectively. Size of nodes (ASVs) according to mean network modularity (*M*, **Extended Data Fig. 6**).

Covariance networks of the rhizosphere microbiomes of control and larva-exposed plants in the four soils showed a significant reduction in mean network modularity in the conducive C10 and C112 rhizospheres when larvae were present, whereas the rhizosphere network in highly suppressive S16 experienced no change (**Fig. 6f, Extended Data Fig. 6**). The S7 rhizosphere presents an intermediate level with a discrete reduction in the mean network modularity in the presence of the larvae (**Fig. 6f**). The decrease in mean modularity coincided with an increase in the number of edges (i.e., connections between ASVs, **Extended Data Fig. 6**).

## Discussion

Here, we tested whether natural suppressiveness of certain field soils against fungal pathogens can extend to insect pests. We first assessed the suppressiveness status of renowned Swiss soils of a 22 km^2^ region near Lake Neuchâtel, known for their long-standing capacity to suppress *T. basicola*-induced black root rot disease in tobacco^11^. We selected six representative soils from the 96 soils originally characterized in the 1980s by Geneviève Défago and coworkers who linked suppressiveness levels to soil geology^21,22,51^. Our findings corroborate their results and document sustained suppressiveness levels against black root rot after decades in all but one of the selected field soils (**Fig. 1b**).

We demonstrate that the innate soil suppressiveness can extend to herbivorous pest insects, specifically the cereal leaf beetle *O. melanopus* (**Fig. 1b**, **Fig. 2b)**. Nonetheless, variations in the level of suppression, specifically soil C10 being more conducive towards the pest insect than to the fungal pathogen, and soil C112 being moderately conducive, suggest that some of the mechanisms involved may be different. The suppressiveness of the Swiss soils studied was previously attributed to the presence of plant-beneficial *Pseudomonas*^11,21^. We confirmed the presence of antifungal and insecticidal *Pseudomonas protegens*^24,28^ in the rhizosphere of plants growing in the most suppressive soil S16 (**Fig. 6d)**, a likely candidate for pathogen suppression in this soil. Conversely, the microbiome of leaf-feeding *O. melanopus* consists of endosymbionts and leaf-endophytic *Pantoea*^57^ whose relative abundances were similar across soils (**Fig. 5**). Nonetheless, differences in *Pantoea* populations in the suppressive S16 soil (**Fig. 5e**) coincide with a lower abundance of *Wolbachia* and other endosymbionts, suggesting that the onset of insect microbiome dysbiosis^58,59^ had started by day 5. *Pseudomonas* was not detected in the insect’s microbiome or the wheat phyllosphere (**Extended Data Fig. 6**), precluding the possibility of soil-plant-insect transmission of insecticidal *Pseudomonas* and suggesting an alternative explanation for the higher pest protection observed in soil S16 (**Fig 2b**).

Phytohormones, particularly jasmonates, are involved in the plant immune response to herbivorous insects^35–37^. We observed a typical jasmonate-mediated response to herbivory, with enhanced OPDA and JA-Ile levels in wheat leaves exposed to *O. melanopus* larvae^60^ (**Fig. 3, Extended Data Fig. 1**). However, plants grown in S16 soil accumulated less herbivory-responsive phytohormones, although this soil was associated with the highest insect mortality (**Fig. 2b**, **Fig. 3bc**). This indicates a reduced stress level in plants growing in S16 soil, despite the insect mortality at day 5 being comparable to those in S7 and C112 soils (**Fig. 2b**). We observed a manifest priming of plants grown in the S16 soil documented by increased plant SA levels in the absence of insects (**Fig. 3d**), which likely explains reduced herbivory stress^61^, and marks the onset of protection against the leaf-feeding insects (**Fig. 2b**). Similarly, BXs are important plant defense compounds against insects^40,44,45^. We observed only a few BXs whose concentration changed with herbivory (**Fig. 4a**). DIMBOA, together with DIMBOA-Glc the primary BX in wheat^44,62^ and effective against insects^40^, had a significantly higher concentration in the leaves of plants growing in the conducive C10 and C112 soils. As with the jasmonates, we observed lower accumulation of DIMBOA in S16 compared to the other soils (**Fig. 4de**). Higher concentrations of the storage form of DIMBOA and HDMBOA correlated positively with larval mortality (**Fig. 4f**), showing a typical plant response^35–37^. However, the active form DIMBOA was negatively correlated with mortality, suggesting that DIMBOA was not the primary agent responsible for the increased larval mortality. Similarly, cyanogenic glucosides were not involved in larval mortality (**Extended Data Fig. 3**) as previously observed with other pest beetles^63^.

We analyzed the microbiome composition of soils, plants and insects to test whether they could have a role in the innate control of the pest *Oulema*. Indeed, the rhizosphere microbiome composition differed between conducive and suppressive soils, with the largest difference between C10 and S16 (**Fig. 6a**). Several plant-beneficial genera associated with plant growth promotion and phytopathogen suppression, *Herbaspirillum, Serratia,* and *Rahnella*^64–68^, were not detectable in the C10 rhizosphere (**Fig. 6e**). We suggest that these genera contribute to the enhanced insect pest resistance in the suppressive soils, strengthened in S16 by the additional presence of plant-beneficial *P. protegens* (**Fig. 6d**). Several mechanisms may contribute, including the priming of plant defenses^36,61^, as observed with SA (**Fig. 3d**). Moreover, covariance analysis of wheat rhizosphere ASVs revealed a destabilizing effect of leaf infestation by *O. melanopus* in the conducive soils, manifesting in a reduced network modularity (**Fig. 6f, Extended Data Fig. 6**). A similar effect has been observed with soilborne pathogens^69,70^, and highlights the importance of the soil microbiome for plant health^59^. Nonetheless, network modularity remained unaltered in the rhizosphere of plants grown in the suppressive S16 soil. This observation is consistent with the dampened response of plants growing in this soil to leaf-feeding *Oulema* (**Fig. 3 and 4**), higher insect mortality (**Fig. 2b**), higher suppression of *T. basicola* disease (**Fig. 1b**), and higher abundance of plant-beneficial bacteria (**Fig. 6de**).

Our work demonstrates that the innate disease suppressiveness of soils towards belowground fungal pathogens extends to above-ground herbivorous pests. This is mediated through a complex soil-plant feedback network involving critical plant-beneficial root-associated bacteria and the priming of plant responses to herbivorous insects (**Extended Data Fig. 7**). The outstanding natural suppressiveness of some soils to both pathogens and herbivorous pests exemplifies the crucial role of soil microbial diversity in plant health and as a reservoir of beneficial bacteria suited to serve as inoculants in environmentally sustainable crop protection strategies.

## Online Methods

### Soil sampling and black root rot suppressiveness assessment

Six field soils were collected in April 2021 in the region of Payerne, Switzerland, based on previous descriptions^11,21–23,51,52^: S16 (46.8849N, 6.9225E), S7 (46.8614N, 6.8983E), S8 (46.8674N, 6.9038E), C10 (46.8634N, 6.9243E), C112 (46.8378N, 6.8877E), and C6 (46.8589N, 6.8841E). The top soil (0-30 cm) was removed to avoid root interference from the cover of mixed plant species. Sixty kg of each soil at a depth of 30-60 cm were collected, manually homogenized, and sieved (mesh size, 10 mm). Soils were tested for suppressiveness against black root rot in tobacco plants caused by *Thielaviopsis basicola* (syn. *Chalara elegans*) as previously described^23^. Briefly, tobacco seeds (*Nicotiana glutinosa* L.) were germinated and seedlings grown for 4 weeks in a plant growth chamber set to 70% relative humidity and 22 °C, with a 16 h light period (880 µmol m^-2^ s^-1^), followed by an 8 h dark period at 18 °C, before being transferred into pots (8x8x8 cm) containing a 1:1 mixture (wt/wt) of the respective field soil and quartz sand (grain size of 0.6-1.6 mm diameter). Seedlings were watered with Knop’s plant nutrient solution^24^. For pathogen inoculation, 5 mL of a suspension 10^5^ endoconidia mL^-1^ in 0.5% Tween 20, prepared from a 4-week-old culture of *T. basicola* strain ETH D127 grown on malt agar (15 g malt extract L^-1^, 17 g agar L^-1^), was inoculated to each plant. Plants were grown for 4 weeks under the same growth chamber conditions as described and the severity of black root rot was assessed using an eight class disease scale of root surface blackened by the presence of chlamydospores of the pathogen, as previously described^21,23^. Ten replicates per condition were performed.

### Cereal leaf beetle plant assay

Adult cereal leaf beetles (*Oulema melanopus*) were collected from wheat fields and reared on barley plants (*Hordeum vulgare* cv. Esprit) growing in oviposition cages placed in a greenhouse chamber set at 70% of relative humidity and approximately 20 °C with a 16 h light period, followed by an 8 h dark period at 16 °C. Pupating larvae were collected and emerging adult beetles that entered diapause were kept at 6 °C for up to three months before placing them onto fresh barley plants, where they mated and laid the eggs used in the wheat infestation experiment described in the following.

Wheat seeds (*Triticum aestivum* cv. Arina) were surface-disinfected in 1.4% NaOCl for 15 min, rinsed with sterile water, and then germinated on 1% agar plates at 22 °C for 48 h. Wheat seedlings were transferred to 50 mL Falcon tubes partly filled with a soil mixture consisting of 5 mL of sterile sandstone grains (grain size of 4 mm diameter), on top of which was added a 1:1 mixture of the respective field soil and sterile quartz sand (grain size of 0.2-4 mm diameter). Plants were watered every 2 to 3 days with 5 mL of sterile distilled H_2_O. The wheat plants were left to grow for approximately 1 week (greenhouse chamber, same conditions as above for barley) until the start of the development of the second leaf. Freshly hatched first instar *O. melanopus* larvae were cleaned on kitchen paper before placing one larva on the first leaf of each plant. The leaves of each individual plant were held to a stick to prevent contact between the plants and the movement of larvae from one plant to the other. The collection and measurements were done at day 5 or 11 , depending on the parameters assessed. Day 5 parameters that were assessed were phytohormones and benzoxazinoids (BXs), as well as plant and insect microbiomes, while day 11 parameters included leaf damage. Larval mortality was evaluated daily until day 11. The experiment was repeated at least two independent times.

### Assessment of insect survival and quantification of leaf damage

Larval survival was assessed daily for 11 days. There were 20 plants per experiment, with one larva per plant. The experiment was run two independent times. *O. melanopus* larvae were considered dead when their surface excrement and mucus cover had dried out and the larvae had fallen off the leaf or did not respond to gentle prodding with a disinfected brush. On day 11, plants were gently removed from the tubes and leaves were scanned. The scanned leaves were used to assess the percentage of leaf damage caused by larval feeding using the Pliman v1.0.0 package in R (https://tiagoolivoto.github.io/paper_pliman/). *O. melanopus* larvae consume the green mesophyll cells between leaf veins but leave the lower epidermis intact, leading to the characteristic transparent longitudinal stripes in the leaves that can be quantified for damage assessment^3^. Example images of damaged and healthy plants were segmented to separate the background from the leaves and used to train the algorithm to 95% accuracy in healthy plants. The remaining samples were analyzed in R.

### Extraction and quantification of phytohormones, benzoxazinoids, and cyanogenic compounds

Leaves were harvested, weighed, and flash-frozen in liquid nitrogen. Roots were first rinsed with distilled H_2_O and gently cleaned before weighing and flash-freezing in liquid nitrogen. Tissue samples from leaves and roots were finely ground while remaining frozen using a mortar and pestle. For phytohormones, approximately 50 mg of powdered tissue was processed as previously described^71^. Measurements were made using a QTRAP 6500 tandem mass spectrometer coupled to an Acquity UPLC I-Class chromatographic system. For extraction of benzoxazinoids and cyanogenic glycosides, eight 2-3 mm glass beads and 1 mL of H_2_O:methanol:formic acid (50:50:0.5, v/v) were added to 25 mg of ground tissues. The samples were homogenized in a bead mill for 3 min at 30 Hz and centrifuged at 12,000 x *g* for 3 min. Two hundred µL of extract was collected per sample, placed in an HPLC vial holding a 250 µL conical insert and analyzed as previously described^39,46,63^.

### Extraction of total DNA from samples

Total DNA from soils, leaves, and rhizospheres was extracted using the DNeasy PowerSoil Pro Kit (Qiagen), while for insects the DNeasy Blood & Tissue Kit (Qiagen) was used. Soil samples were prepared by adding 10 mL of minimal medium (MM^72^) to 10 g of soil. For rhizosphere samples, roots were gently shaken to remove excess soil, pooled in groups of three, and placed in tubes with 10 mL of MM. The samples were vortexed for 20 min and left to stand for 3 min to allow large particles to sediment. The supernatant was then centrifuged at 8,000 x *g* for 1 min to pellet cells, which were stored at -20 °C until DNA extraction.

Leaf samples from individual plants, weighing approximately 130 mg, were cut from the plant, placed in 2-mL sterile tubes and stored at -20°C. Ten sterile glass beads (a mix of 0.75-1 mm and 3-5 mm beads) were added per sample with 500 µL of sterile PBS (pH 7.4). The tubes were then homogenized for 30 seconds at 60 Hz in a bead mill homogenizer and centrifuged for 1 min at 300 x *g*. The resulting supernatant was transferred to the PowerSoil kit (Qiagen) for DNA extraction according to the manufacturer’s instructions.

For DNA extraction from *O. melanopus*, individual larvae were cleaned by first removing the protective fecal and mucus cover with a paper towel. The larvae were then surface disinfected by rinsing in 70% ethanol for 20 s, followed by washing in 0.05% SDS, and rinsing again for another 20 s in 70% ethanol. Finally, insects were rinsed in sterile distilled H_2_O for 20 s before being dried on filter paper and stored at -20°C. Five larvae were pooled together and macerated following the same bead mill homogenizer steps described above. The resulting supernatant was transferred to the Blood & Tissue kit (Invitrogen) for DNA extraction according to the manufacturer’s instructions.

The obtained DNA was kept at -20 °C until further processing. DNA concentration was measured using the Qubit dsDNA HS Assay kit (Invitrogen).

### Amplicon sequencing of the 16S rRNA gene and analysis

To prepare samples for sequencing, 10 ng of DNA per sample were used to amplify the V3-V4 region of the 16S rRNA gene using the 341F (5′-CCTACGGGNGGCWGCAG-3′) and the 805R (5′-GACTACHVGGGTATCTAATCC-3′) primers^73^, following the Illumina 16S Metagenomic Sequencing Library preparation protocol as previously described^72^. Samples were sequenced at the Lausanne Genomic Technologies facility (Lausanne, Switzerland), using an Illumina MiSeq v3 instrument running for 300 cycles. Raw sequences were filtered and trimmed by quality using fastp v0.32.2^74^, and further processed following the DADA2 v1.20.0 pipeline^75^, as previously described^72^, until the obtention of amplicon sequence variants (ASVs). Taxonomy was assigned using the SILVA v138 database^76^. ASV sequences, taxonomy and metadata were imported into the phyloseq v1.36.0 R package^77^ for diversity and compositional analyses as previously described^72^. Association between ASVs was assessed using the sparse inverse covariance estimation for ecological association inference (SPIEC-EASI), SpiecEasi v1.1.2 R package^78^, using the top 250 most abundant ASVs as previously described^72^, and performing 18 independent repetitions per network.

### Statistical analyses

The normality of the data was tested using Shapiro’s and Levene’s tests. Differences were assessed using the non-parametric Kruskal-Wallis rank-sum test within the agricolae v1.3-5 R package^79^. Post hoc analyses were performed using Fisher’s least significant difference (LSD). The *P* values were corrected using the false discovery rate (FDR). Different statistical groups were defined at a *P* ≤ 0.05. To assess fold changes, the individual values of the insect treatment groups were divided by the corresponding mean concentration of each control group. This was then log2-transformed. Insect survival data was analyzed using the R packages survminer 0.4.9, survival 3.3-5, multcomp 1.4-25, and coxme 2.2-18.1, using the randomization effect by experimental run for the mixed-effect Cox regression. The effect of variables on Bray-Curtis microbiome dissimilarities was evaluated using permutational multivariate analysis of variance (PERMANOVA, 999 permutations) with the *adonis2* function of the vegan v2.6-4 R package^80^. Correlations between BXs concentrations and *O. melanopus* larval mortality were assessed using the Spearman correlation of the *stat_cor* function within the ggpubr R package.

## Acknowledgements

We thank Caterina Matasci (Delley Seeds and Plants Ltd., Switzerland) and Gernot Alber (SOTA, Payerne, Switzerland) for providing the wheat and tobacco seeds, respectively, used in this study. We thank the Lausanne Genomic Technologies Facility in Lausanne, Switzerland, for sequencing the samples, and the *Serre* platform of FR3728 BioEnviS at Université Lyon 1 for plant disease trials. We acknowledge and thank Irena Todorovic, Lorenzo Pepe, Maëlle Corminboeuf and Océane Devisme for their assistance in experimental setup and sample collection. We acknowledge Claudia di Cesare and Sylvie Guinchard from the Neuchâtel Platform of Analytical Chemistry for their assistance in plant metabolite extraction and processing.

## Author contributions

Conceived and designed the experiments: PV, YML, TS, CK and DGS Performed the experiments: NH, PV, GG, FK, CMH, AA, JV, DM and DGS Analyzed the data: NH, PV and DGS Contributed materials/analysis tools: GG, DM, YML, TS, CK and DGS Wrote the paper: NH, CK and DGS

## Data availability

All data generated in this work together with the scripts used for their analyses have been provided throughout the text, in the supplementary files, or are publicly available in GitHub (https://github.com/nhrmsn/SuppressSoil-Data). Raw read sequences of the 16S rRNA gene amplicons have been deposited in the NCBI Read Sequence Archive database and are publicly available under the BioProject accession number PRJNA1075215.

## Extended Data Figures

**Extended Data Fig. 1.**
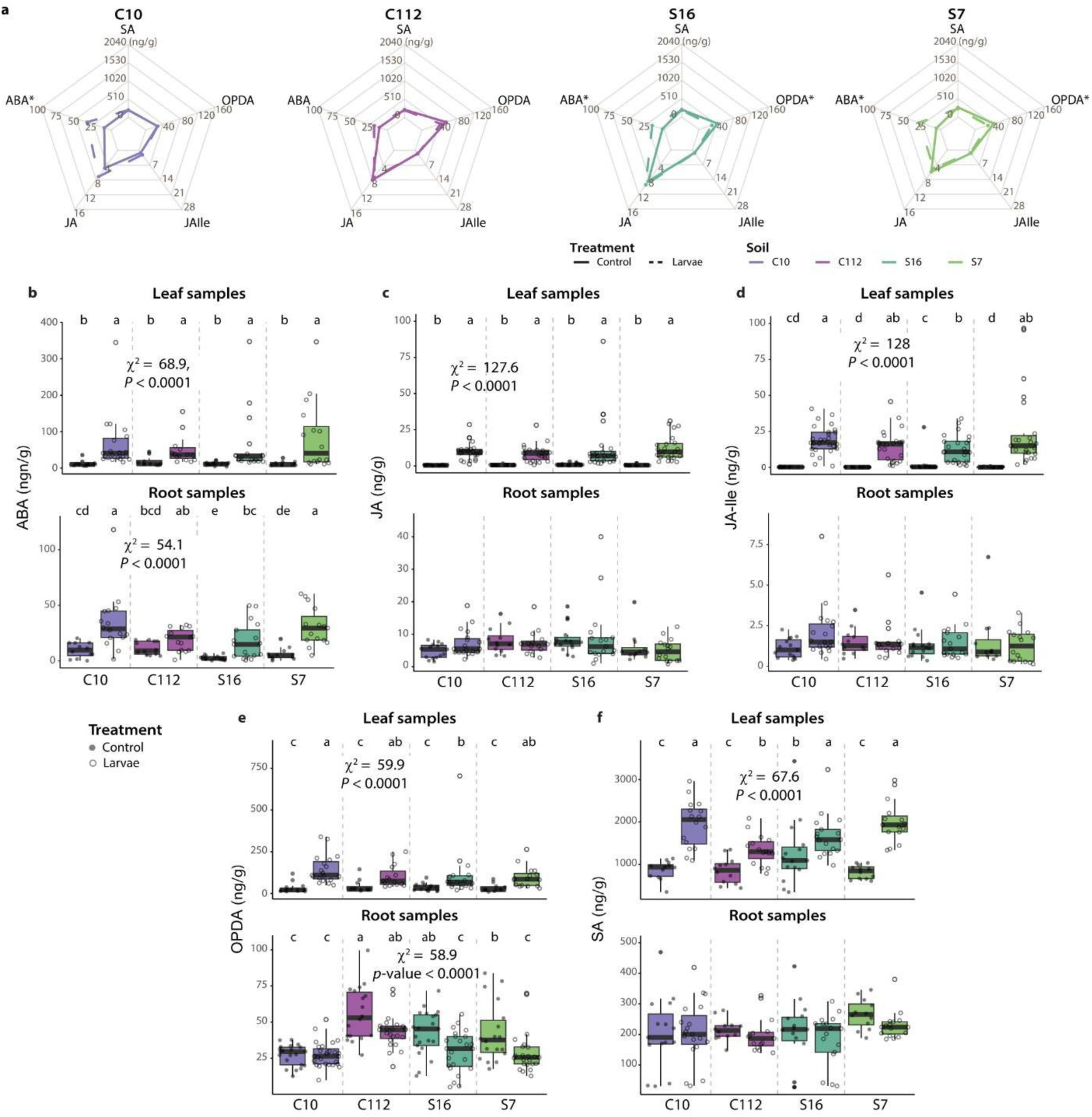
Changes in levels of defense-related phytohormones in wheat roots and leaves exposed to *Oulema melanopus* larvae. **a**, Spider plots show insect and control group concentrations of abscisic acid (ABA), jasmonic acid (JA), jasmonyl-isoleucine (JA-Ile), 12-oxo-phytodienoic acid (OPDA), and salicylic acid (SA) in root samples in the soils C10, C112, S16, and S7. Asterisks denote phytohormones whose concentrations differed significantly between insect and control groups (Kruskal-Wallis, *P* ≤ 0.05). **b-f**, Boxplots representing the concentration of (**b**) ABA, (**b**) JA, (**d**) JA-Ile, (**e**) OPDA, and (**f**) SA in plants that had been exposed or not to the insect in the four soils (*n* ≥ 12, at least two experimental runs were performed). The points represent individual replicates. Statistical differences were assessed using the Kruskal-Wallis test and LSD post hoc analysis. The *P* values were corrected by FDR. Different letters indicate significant differences between groups (*P* ≤ 0.05).

**Extended Data Fig. 2.**
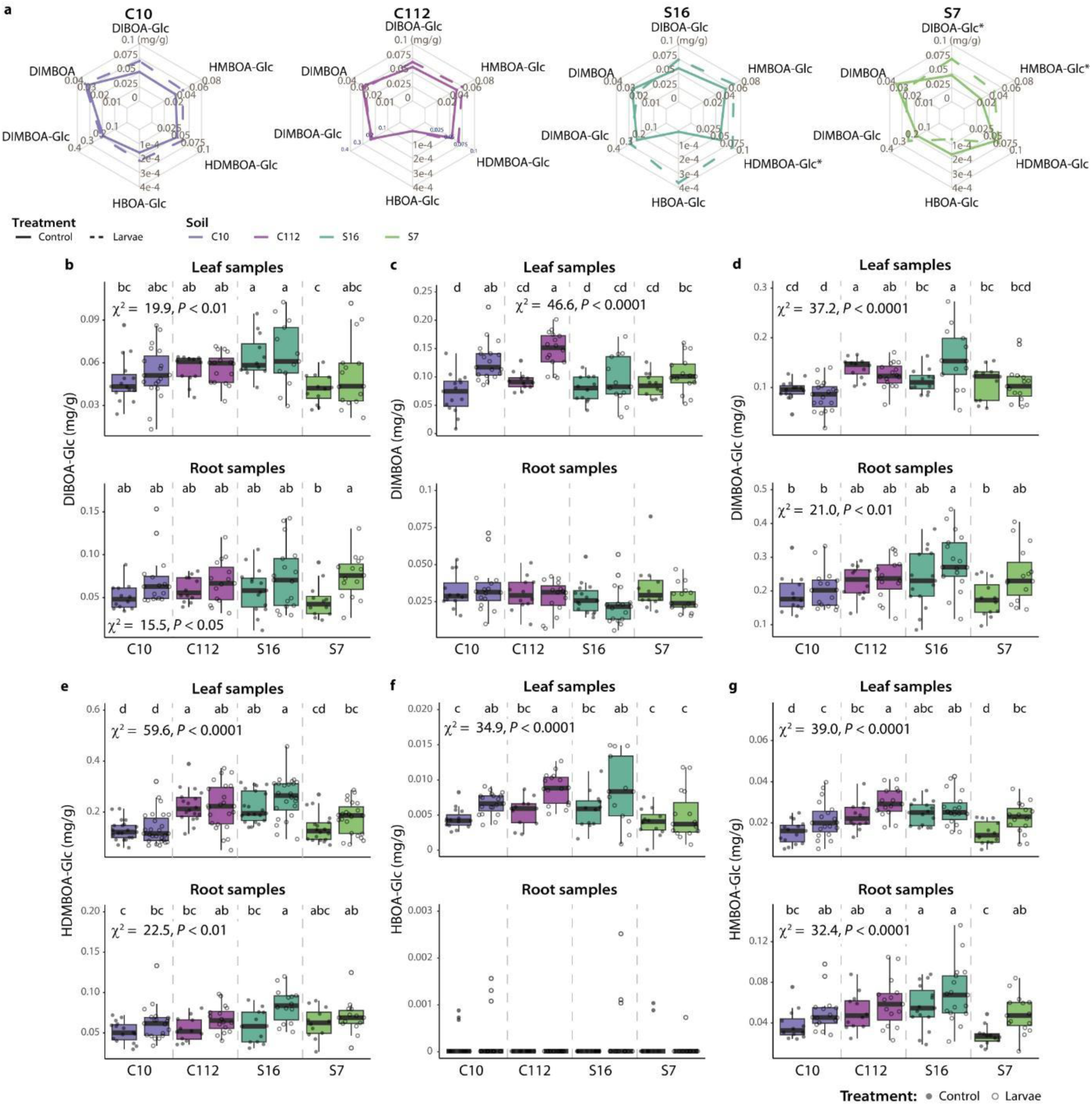
Changes in benzoxazinoid (BX) concentration in wheat roots and leaves exposed to *Oulema melanopus* larvae. **a**, Spider plots show *O. melanopus* larvae and control group concentrations of DIBOA-Glc, DIMBOA, DIMBOA-Glc, HDMBOA-Glc, HBOA-Glc, and HMBOA-Glc in root samples in the soils C10, C112, S16, and S7. Asterisks denote BXs whose concentrations differed significantly between insect and control groups (Kruskal-Wallis, *P* ≤ 0.05). **B-g**, Boxplots representing the concentrations of (**b**) DIBOA-Glc, (**c**) DIMBOA, (**d**) DIMBOA-Glc, (**e**) HDMBOA-Glc, (**f**) HBOA-Glc and (**g**) HMBOA-Glc in root or leaf samples exposed or not to the larvae in the four soils (*n* ≥ 12, two experimental runs for leaf samples, one for root samples). The points represent individual replicates. Statistical differences were assessed using the Kruskal-Wallis test and LSD post hoc analysis. The *P* values were corrected by FDR. Different letters indicate significant differences between groups (*P* ≤ 0.05).

**Extended Data Fig. 3.**
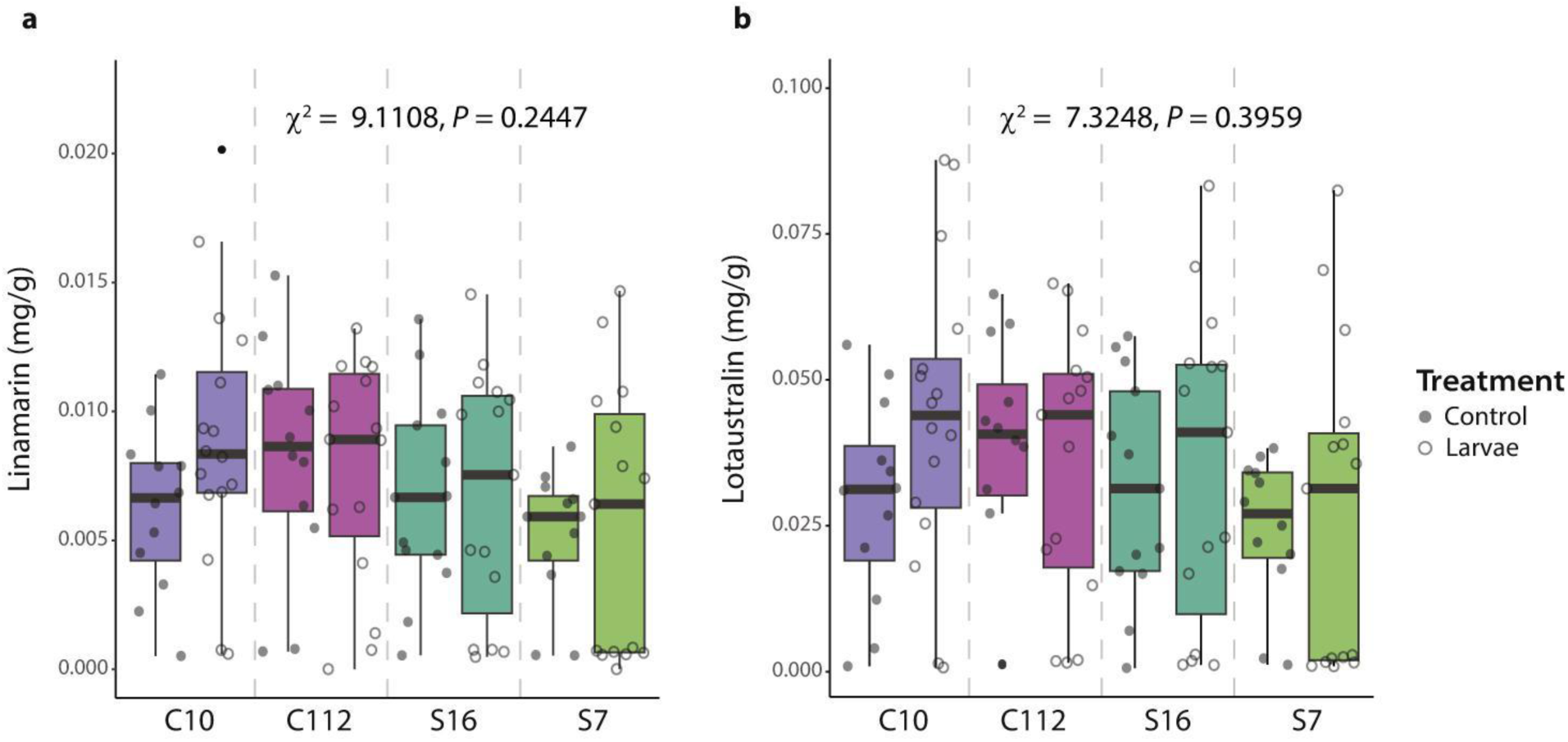
Concentration of the cyanogenic compounds linamarin and lotaustralin in leaves of plants exposed to *Oulema melanopus* larvae. **a,b**, Differences in the concentrations of (**a**) linamarin and (**b**) lotaustralin between plants growing with *O. melanopus* larvae and control plants (*n* ≥ 12, one experimental run). Box plots showing the concentrations in mg per gram of tissue. The points represent individual replicates. There were no statistical differences (*P* ≥ 0.05) between samples based on the Kruskal-Wallis test.

**Extended Data Fig. 4.**
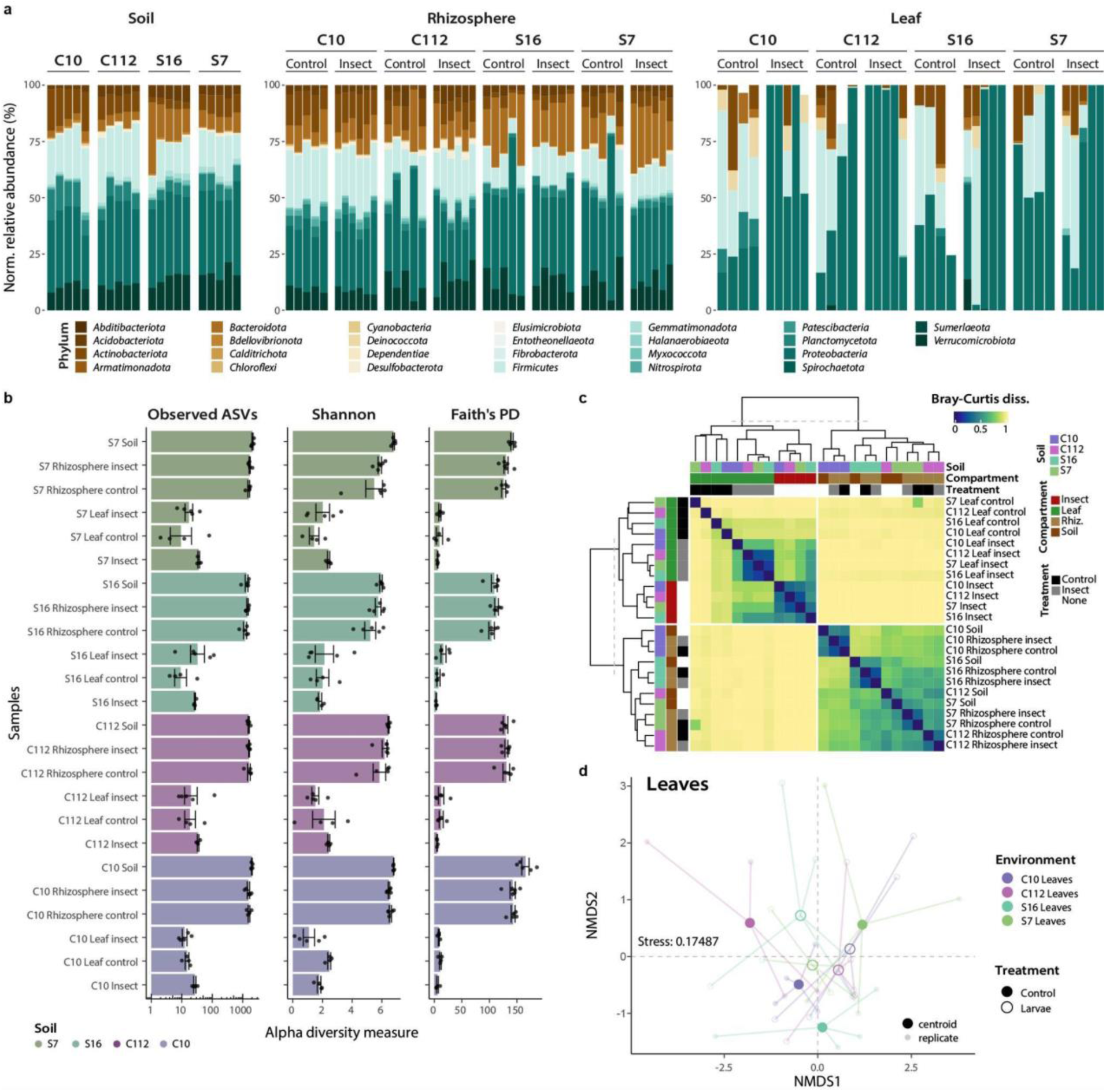
Microbiome composition and diversity of soils, wheat rhizospheres/phyllospheres and *Oulema melanopus* insects. **a**, Cumulative sum scaling (CSS)-normalized relative abundance of ASVs in samples at the phylum level. **b**, Alpha diversity measurements (observed ASVs, Shannon diversity and Faith’s phylogenetic diversity (PD)) across samples. Bars represent mean values ± s.e. (*n* ≥ 4). Black dots represent individual replicates. **c**, Complete-linkage clustering of samples based on Bray-Curtis dissimilarities and heatmap. Replicates were merged using the mean ASVs abundance. **d**, Non-metric multidimensional scaling (NMDS) ordination analysis of leaf samples based on Bray-Curtis dissimilarities.

**Extended Data Fig. 5.**
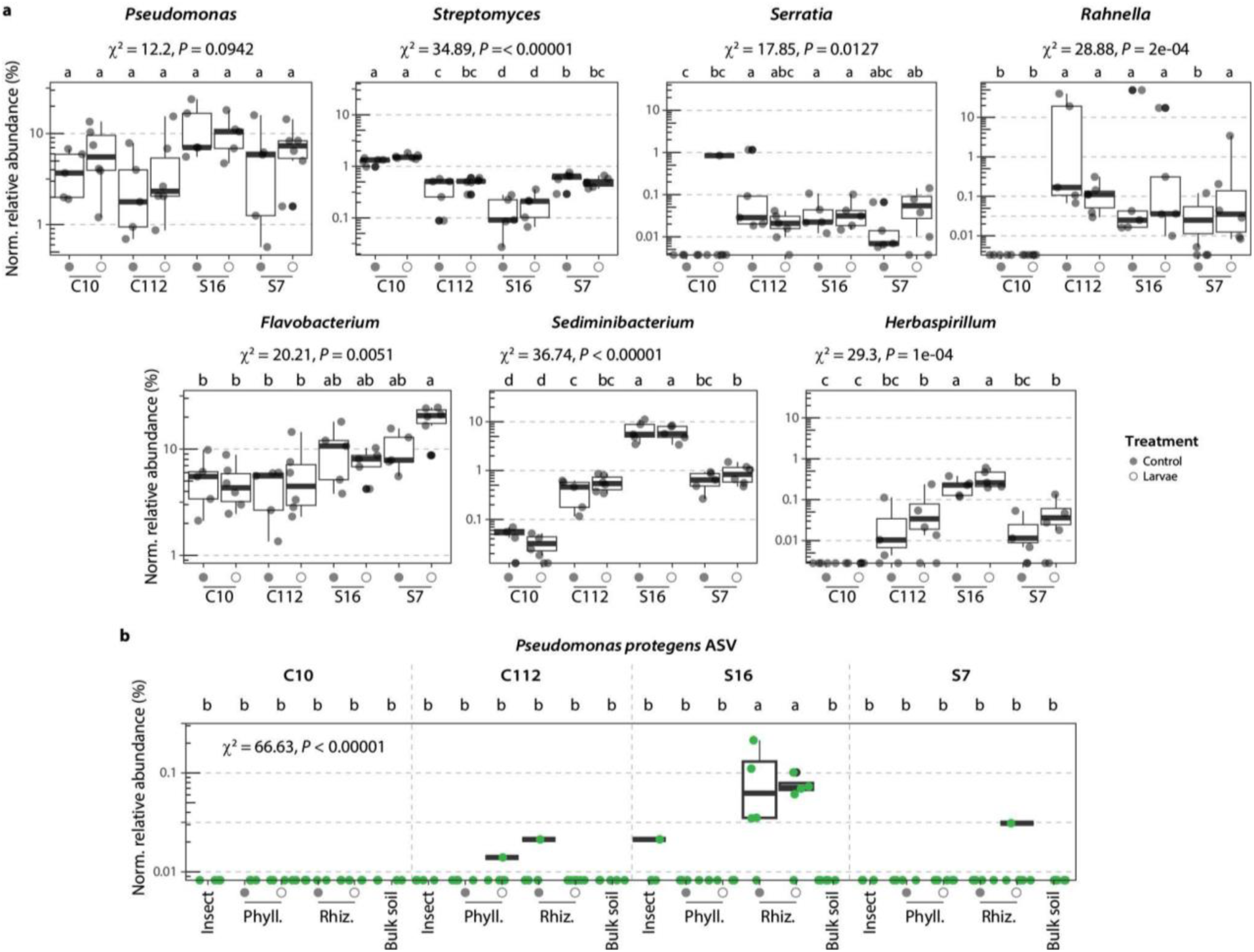
Relative abundance of key bacterial taxa throughout samples. **a**, CSS-normalized (Norm.) relative abundance of genera showing differential abundance changes in the rhizosphere of wheat plants growing in the different soils exposed or not to *Oulema melanopus* larvae. **b**, Norm. relative abundance of *Pseudomonas protegens* CHA0^T^ (exact ASV sequence match, green dots) across microbiomes: Insect (*O melanopus* larvae), wheat phyllosphere (Phyll.), rhizosphere (Rhiz.) or the bulk soil. Dots represent individual replicates (*n* ≥ 4). Dots collapsed at the bottom of the plots signify absence of the taxa (i.e., relative abundance = 0). Significant differences were assessed using the Kruskal-Wallis rank sum test and post hoc analysis by means of the Fisher’s least significant difference. The *P* values were corrected by FDR. Different letters indicate significant differences between groups (*P* ≤ 0.05)

**Extended Data Fig. 6.**
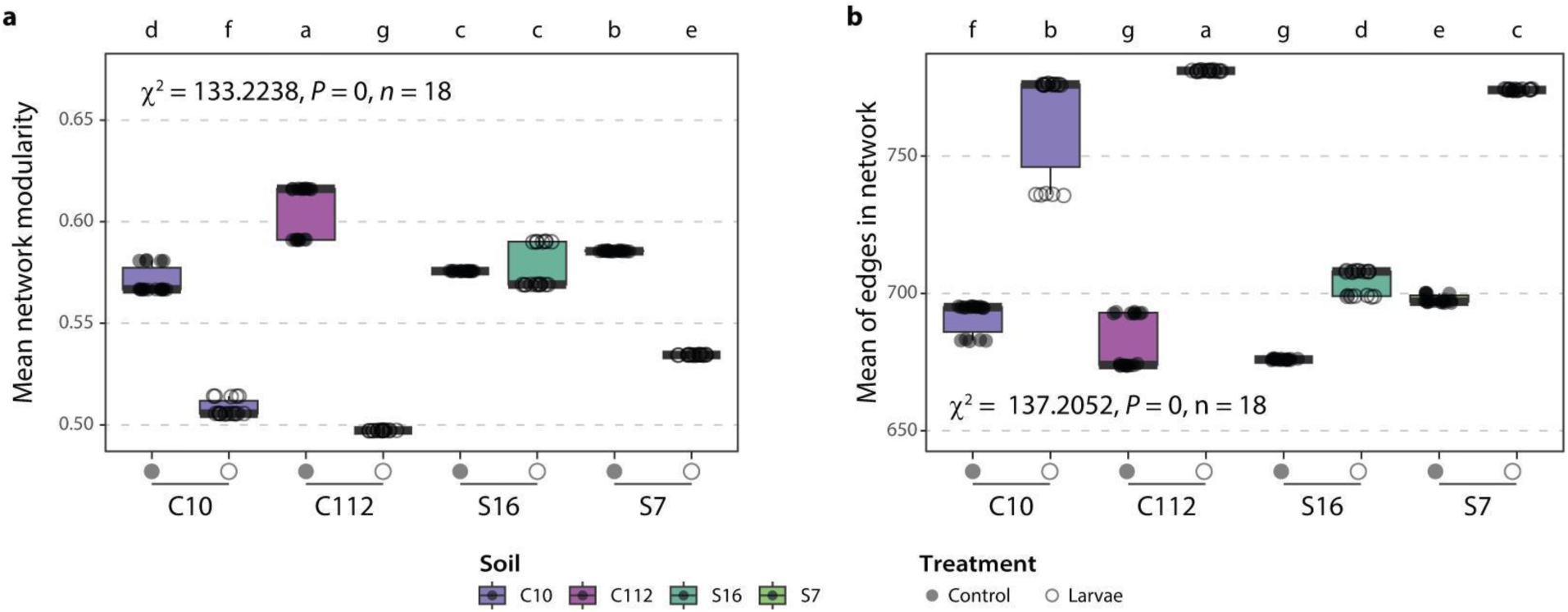
Network modularity and number of edges in the rhizosphere of wheat plants exposed or not to *Oulema melanopus* larvae. **a,b**, Mean network modularity (**a**) or mean number of edges (**b**) in association networks based on sparse inverse covariance estimation among the top 250 ASVs from rhizosphere microbiome samples in control plants or those exposed to *O. melanopus* larvae herbivory across the four soils studied. Statistical differences were assessed using the Kruskal-Wallis test and post hoc analysis using Fisher’s least significant difference. The *P* values were corrected by FDR. Different letters indicate significant differences between groups (*P* ≤ 0.05).

**Extended Data Fig. 7.**
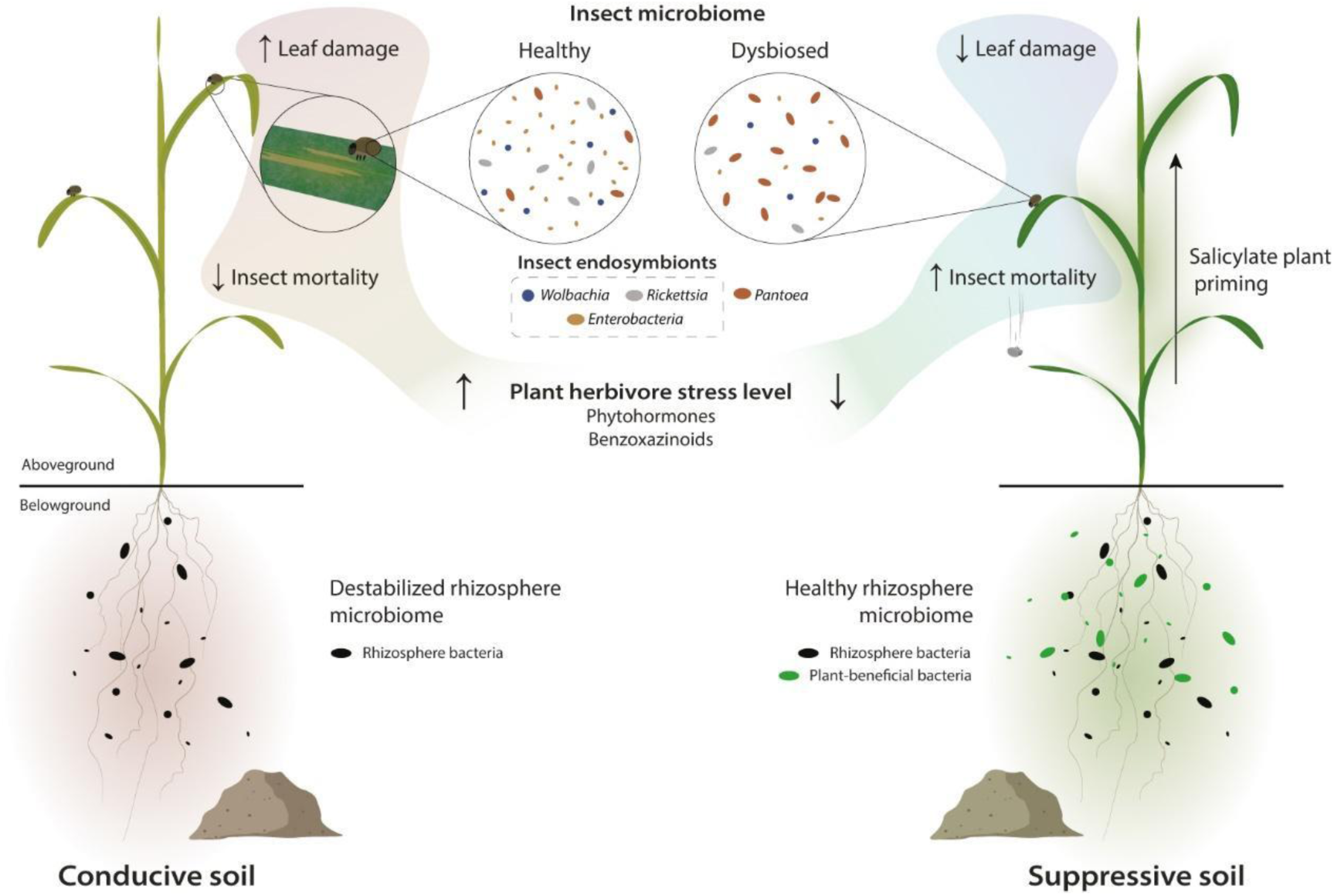
Proposed model for innate soil suppressiveness towards herbivorous insects. Plants grown in natural conducive (disease-favoring) soils (left) show typical aboveground plant-mediated responses to herbivorous insects, manifesting in increased levels of herbivore defense-related phytohormones and benzoxazinoids. Belowground, the root-associated microbiome becomes dysbiosed when the pest insect feeds on the leaves. Conversely, plants grown in suppressive soils (right) have plant-beneficial bacteria associated with their roots. In addition, a higher level of salicylate operates in leaves of suppressive soil grown-plants, which makes them become less stressed and prepared for a faster response when exposed to the insect and thus, insect defense-related phytohormones and benzoxazinoids have a lower response. The displacement of endosymbionts in the microbiomes of insects feeding on plants growing in suppressive soils reflects the onset of a higher insect mortality. This intricate innate soil-plant feedback network ultimately protects plants from the herbivorous pest.

## Supplementary material

The following supporting information is available for this article:

**Supplementary Note 1**

**Supplementary Note 2**

### Supplementary Note 1

The *Oulema melanopus* mortality assay shows that larvae feeding on plants growing in soil S16 had the highest mortality rate, with all the larvae dead by day nine, while for soils S7 and C112 >25% of the larvae were alive at the end of the experiment (**Fig. 2b**). Larvae feeding on plants growing in the soil C10 showed the highest survival rate by day 11, but still 50% of the larvae died. This seemingly high mortality rate, even in the most conducive soil C10, coincides with normal population dynamics of *O. melanopus* larvae observed previously^1,2^. Despite this, all mortality experiments were conducted under the same conditions, and there were significant differences in larval mortality depending on the soil in which the plants were grown (Cox model, *P* = 0.0001035, *n* = 20, two experimental runs per soil).

### Supplementary Note 2

Due to its higher stability and the analytical method employed, only the aglucone form of DIMBOA has been measured. To comply with space constraints, the full chemical IUPAC names of the benzoxazinoids studied have been omitted from the main text, and only the abbreviated name is provided. Their full IUPAC names are as follow: DIBOA-Glc: 4-hydroxy-2-[3,4,5-trihydroxy-6-(hydroxymethyl)oxan-2-yl]oxy-1,4-benzoxazin-3-one, DIMBOA: 2,4-dihydroxy-7-methoxy-1,4-benzoxazin-3-one, DIMBOA-Glc: 4-hydroxy-7-methoxy-2-[3,4,5-trihydroxy-6-(hydroxymethyl)oxan-2-yl]oxy-1,4-benzoxazin-3-one, HBOA-Glc: 2-[3,4,5-trihydroxy-6-(hydroxymethyl)oxan-2-yl]oxy-4H-1,4-benzoxazin-3-one, HDMBOA-Glc: 4,7-dimethoxy-2-[3,4,5-trihydroxy-6-(hydroxymethyl)oxan-2-yl]oxy-1,4-benzoxazin-3-one, HMBOA-Glc: 7-methoxy-2-[3,4,5-trihydroxy-6-(hydroxymethyl)oxan-2-yl]oxy-4H-1,4-benzoxazin-3-one.

